# Sinking diatom aggregates provide carbon to drive microscale denitrification in a bulk oxygenated ocean

**DOI:** 10.1101/2022.09.26.509232

**Authors:** Davide Ciccarese, Omar Tantawi, Irene Zhang, Desiree Plata, Andrew R. Babbin

## Abstract

Sinking marine particles drive the biological gravitational pump that naturally sequesters carbon dioxide from the atmosphere. Ubiquitous throughout the ocean, these particles are largely composed of phytoplankton that aggregate together or are repackaged by zooplankton into pellets that sink to the deep. Despite their small size, the compartmentalized nature of these particles promotes intense localized metabolic activity by the bacteria lucky enough to colonize them. Due to their sheer numbers, these microscale interactions can change the chemistry of the bulk ocean and impact global biogeochemical budgets. As soon as phytoplankton-derived particles are exported from the surface ocean, the fate of the carbon depends on the lability and availability of the carbon, the diffusive supply of oxidants from the bulk, and the development of microbial communities throughout the aggregate. Here we show with a model experimental system that aggregates composed of marine diatoms — important primary producers substantially contributing to global carbon export — can support active denitrification even among bulk oxygenated water ill-conducive to anaerobic metabolisms. We further show the primary nitrite maximum could be formed, in part, due to dissimilatory reduction of nitrate and nitrite occurring at anoxic microsites within such particles. Particle-based denitrification and other anaerobic metabolisms can change the global budget of elemental cycles important for life and climate across the oceans.

## MAIN TEXT / INTRODUCTION

The microbial metabolism of denitrification exerts a principal control on the ocean’s global productivity as fixed, bio-available nitrogen limits primary production across most of the global ocean. This anaerobic pathway conducted by specialized organisms is characterized by the stepwise reduction of nitrate through nitrite, nitric oxide, and nitrous oxide to dinitrogen gas. Denitrification occurs when dissolved oxygen concentrations are sufficiently low as to minimize the oxygen poisoning of the enzymes themselves and the organisms facultatively switch from solely respiring aerobically ^1^. Because of the dependence on limited oxygen availability, marine denitrification has classically been investigated in sediments and the ocean’s oxygen deficient zones (ODZs) of the eastern tropical Pacific and Arabian Sea ^2^. Further, global biogeochemical models typically only consider these two marine settings when closing the fixed nitrogen budget ^3^.

Sinking marine particles, however, have been largely excluded from consideration as potential denitrification hotspots of global significance. This marine snow combines the essential features that give rise to denitrification in sediments and ODZs: physical confinement acts to exert a diffusion limit of oxygen supply from the oxygenated bulk and particles are locally enriched in rich organic matter. These two factors combine to make marine particles likely candidates for hosting denitrification ^4–7^, but unlike ODZs and sediments, one distributed across the global water column. Real marine particles, to date, are difficult to sample intact and in sufficient quantities to measure metabolites, chemistry, or microbial activity because of their ephemeral and dispersive nature throughout the ocean. Here we developed and utilized a laboratory system that replicates marine snow in the ocean, composed of marine organic matter that moves through the water column. We show the ease with which oxygen concentrations are reduced locally in a particle, determine where denitrification enzymes are turned on and activity distributed within a particle, and elucidate how microscale dynamics influence the bulk water column chemistry.

To approximate in-situ marine snow particles, we suspended a culture of the marine diatom *Chaetoceros affinis*, a globally important phytoplankton found in productive waters ^8–11^ in agarose (Fig.1, a, b). The agarose here is minimally bio-available yet acts to stabilize the particle and keep it together, similar to various transparent exopolymer (polysaccharide) particles (TEP) in the ocean ^12–16^. *C. affinis* produces much TEP, particularly for its size ^17^. The culture was mixed with an assemblage of organisms isolated from the phycosphere itself, each shown to have facultatively anaerobic growth potential. Suspending the mixed cultures in agarose in such a fashion replicates the way diatomaceous marine snow would form, i.e., the clumping of cells with TEP ^12,18–21^ and their representative phycospheres ^22–26^ being distributed throughout. We lastly suspended oxygen-sensitive nanoparticles in the aggregates to track precisely the microscale dynamic consumption of oxygen throughout the aggregates (Fig. 1 b) and the local consumption within each expanding individual bacterial colony (Fig. 1 c). The agarose discs were sandwiched between two impermeable glass plates and seawater was flown around them at a speed of 4.8 m d^−1^, similar to sinking speeds in the ocean of loose diatom flocculated particles ^18,27–29^. The particles generated, 1.5 mm in radius, were imaged every hour for over 10 days with epifluorescence microscope and the seawater flowing around the particles was sampled once daily for bulk chemistry (nitrate, nitrite, and total dissolved organic carbon).

**Figure 1.**
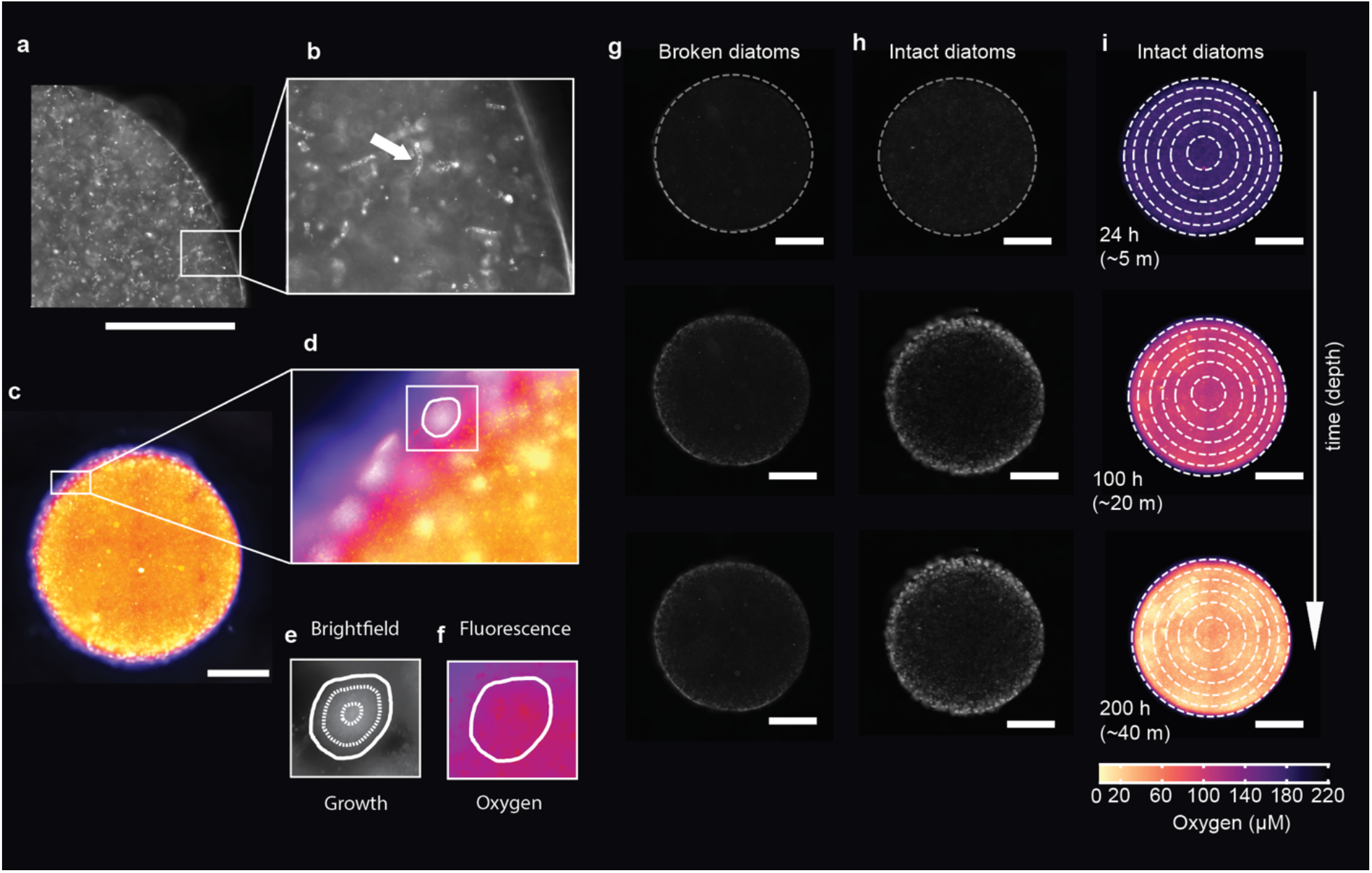
Particle seeded with diatoms and their oxygen evolution during sinking. **a)** Microscopy image of a particle displaying the intact diatoms seeded within the hydrogel and the resulting bacterial colony development. **b)** Close up of intact diatoms within the hydrogel particle. A white arrow indicates an intact diatom chain. **c)** Pseudo colored particle displaying the change in oxygen concentration across the particle as measured by fluorescent nano-reporting particles. **d)** Close up of the edge of a particle showing a steep gradient of oxygen at the edge. One colony is highlighted with a white contour. **e)** The area of each colony is measured at every time point, using brightfield images. The dashed lines denote the 24, 100, and 200 h time points. **f)** The oxygen consumed within each colony is measured by quantified the fluorescence of nanosensor encased within the microcolony’s biomass. **g** and **h)** Particles seeded with broken diatoms and intact diatoms, respectively. Visualization of the bacterial biomass accumulation at three depths 5 m, 20 m and 40 m. **i)** Increase of nanosensor fluorescence as the oxygen is consumed over the sinking depth. The white scale bar in all panels equals 1000 μm.

In the real ocean, there are two dominant pathways of marine snow generation: the consumption and repackaging of phytoplankton into fecal pellets by zooplankton ^30,31^ and the spontaneous clumping of aggregates, which particularly is enabled during blooms ^31–33^. We explored both conditions, by seeding our experiments with broken diatoms and intact diatoms, respectively. Each represents an endmember of the spectrum of carbon availability dictated by the intactness of the diatom cells, with the former releasing all the dissolved carbon rapidly in a pulse and the latter maintaining more carbon for longer via a slow leak. We hypothesize that the broken diatoms should induce a *boom-and-bust* scenario for facultative copiotrophs able to respond rapidly to the nutrient pulse whereas the intact cells should permit a slower but ultimately more abundant community to develop. Moreover, the oxygen drawn down inside the particle should be more intense from intact cells because the organic nutrients are better retained. Combined, we anticipated that the intact diatoms would lead to less dissolved organic carbon (DOC) leaked to the water column than the broken diatoms. Yet, how readily denitrification can rise within this system will depend on the speed of microbial growth on natural phytoplankton exudates compared to the flushing rate by diffusion at the scale of the bacterial microcolonies themselves. Due to the ephemeral nature of sinking particles and their optical inaccessibility, the dynamics of how microscale processes impact the macroscale remain largely unexplored empirically. The novel method employed here overcomes these limitations and opens a window into the highly spatiotemporal dependent scale of heterogeneous biomass development, microscale oxygen consumption, and bulk chemistry evolution.

## RESULTS AND DISCUSSION

### Selective pressures within the phycosphere

To generate an inoculum with which to seed our particle experiments, we first had to isolate organisms from the milieu of the phycosphere ^34^. We easily isolated a number of bacterial members of the community, many of which tested positive for the ability to reduce nitrate and nitrite. In all, we seeded our particles with a mixed assemblage of five bacterial species, each representative of different genera: *Alteromonas, Marinobacter, Phaeobacter, Idiomarina*, and *Virgibacillus*. The ease with which facultative denitrifiers can be found in the phycosphere emphasizes the potential of localized denitrification in phytoplankton-derived marine snow. As a carbon source, we used the chain-forming diatom *Chaetoceros affinis*, an organism of ecological significance, and one that produces extracellular polysaccharides and releases carbohydrates and amino acids throughout its lifecycle ^11^. This organism has also been shown to attract bacteria to cultivate its own phycosphere ^25^.

When diatom-bacterial aggregates form in the water column, interactions at the microscale unfold rapidly. The results of these interactions impact the efflux of phytoplankton exudates and metabolic products that ultimately influence water column biogeochemistry. The diatoms leak their organic contents that serve as nutrients for bacterial heterotrophs across an aggregate. These diverse bacteria in turn can interact to form self-engineered compartmentalized spaces that optimize the supply of necessary nutrients via diffusion. Yet, if bacteria cannot grow fast enough to intercept the labile carbon leaking from the phytoplankton around them, the dissolved organic carbon resource is lost from the aggregate environment. To understand how local gradients drive bacterial productivity across space and its link to the larger scale processes, we quantify the biomass accumulation with respect to the distance to edge of a particle (Fig. 2). Increased growth manifests closer to the edge than the center in both the intact and broken diatom conditions (Fig. 2 a, e) because even though the source of organic carbon is distributed throughout the particle, the oxidants (oxygen and nitrate) are resupplied from the bulk. As such, it is the oxidants that limit the colony development. Yet, the integrity of the diatoms further plays a role as the broken diatoms do not permit as much growth as their intact counterparts (Fig. 2 b, f). This is likely because the initial organic reserves contained with the cells are released too quickly, resulting in their ultimate loss to the bulk seawater rather than their bacterial consumption and recycling.

The distributions of biomass at the end of the experiments in both broken and intact diatom particles show a comparable decay across the particles (Fig. 2 c, g), with the colonies closest to the edge being larger. Yet, the intact diatoms supported much more growth of the colonies closest to the edge than the broken diatoms did. The number of colonies in particles with broken diatoms is slightly higher in number closer to the edge, compared to number of colonies that emerge in the particles seeded which their distribution is more evenly distributed across the entire particle (Fig. 2 d, h). The larger colonies supported by the intact diatoms do not permit as numerous colonies to develop because of competition.

**Figure 2.**
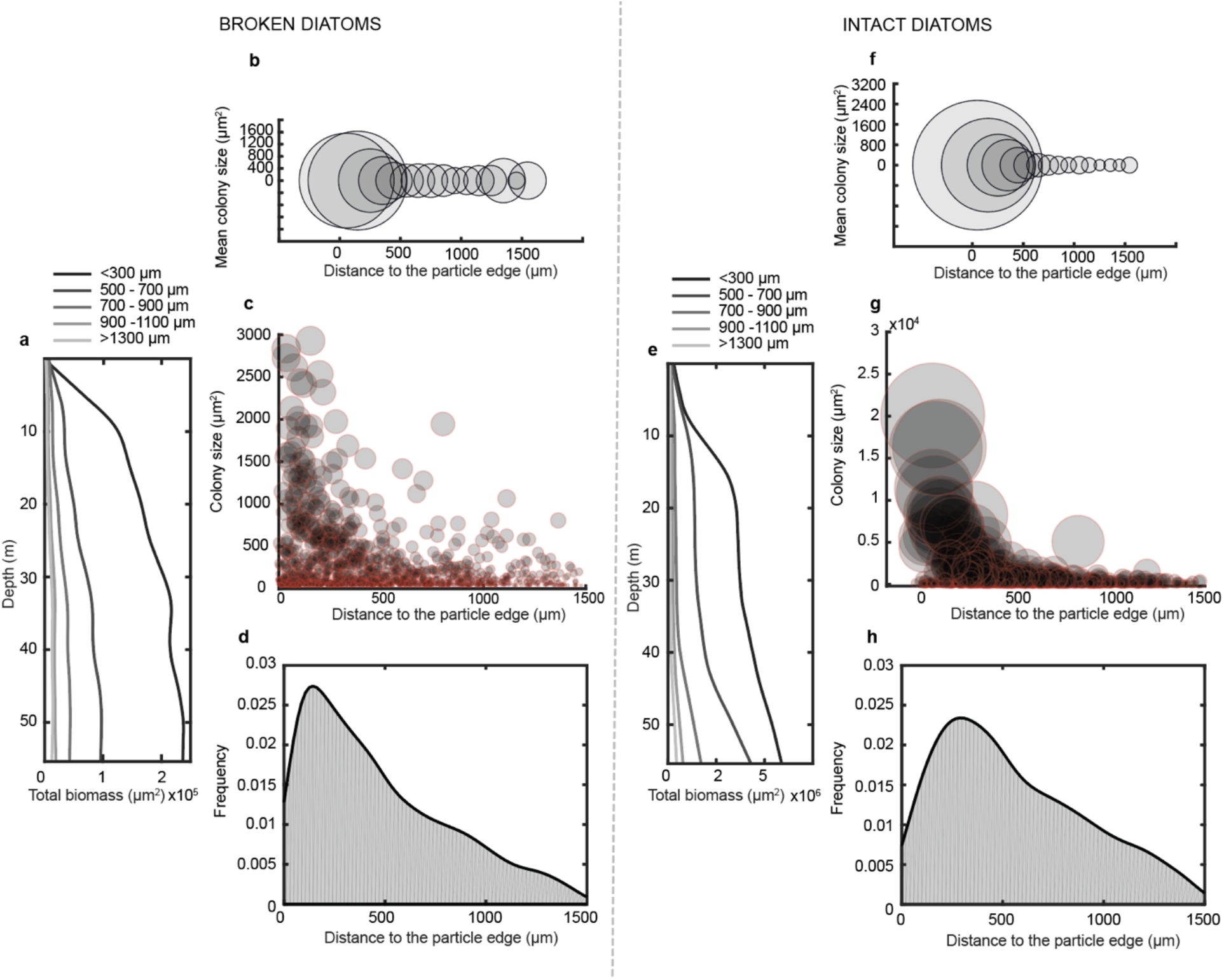
Broken and intact diatoms drives a spatially heterogeneous biomass accumulation. **a)** Translated depth profile of the cumulative biomass productivity at different distances from the edge of particles seeded with broken diatoms (n = 4). **b)** Mean colony size of particles seeded with broken diatoms (n = 4) at different locations within the particles. **c)** Decay of colony size from the edge of particles seeded with broken diatoms (n = 4), each dot size is proportional to the size of the colony. **d)** Probability density function of the distribution of microcolonies along the radius of particles seeded with broken diatoms (n = 4). **e–h)** The same as a–d but using intact diatoms instead of broken ones.

By monitoring the dissolved organic carbon that escapes from the particle into the bulk seawater, we can relate biomass growth dynamics with the bulk carbon retained within a particle that is not lost to the flowing bulk environment. The particles seeded with broken diatoms (Fig. 3 a) initially leak a higher (two-sample t-test, p = 0.003) amount of carbon (28.0 ± 0.7 mg of C/L mean ± stdev) compared to the particles seeded with intact diatoms (14 ± 3.6 mg of C/L mean ± stdev). More DOC escapes the particle initially, i.e., the analogue of the export horizon, thus indicating less organic carbon availability maintained locally within the particle to support bacterial biomass accumulation. Yet, notably, particles seeded with intact diatoms show less release of DOC to the bulk (Fig. 3 b), maintaining similar effluxes across the entire lifetime of the particle, retaining more carbon, and leading to greater ultimate bacterial biomass. The impact of these two different modes of organic leakage can be profound: the leakier broken cells will lead to less export of organic matter via the gravitational pump and greater remineralization in shallower waters. This in turn will limit the sequestration of carbon into the deep ocean. Conversely, intact phytoplankton not only retain their carbon better, but the bacteria that comprise their phycosphere are able to recycle a larger fraction of the carbon that leaks and maintain it in the sinking particle, permitting higher export fluxes to deeper depth and higher magnitudes of sequestration for longer times.

**Figure 3.**
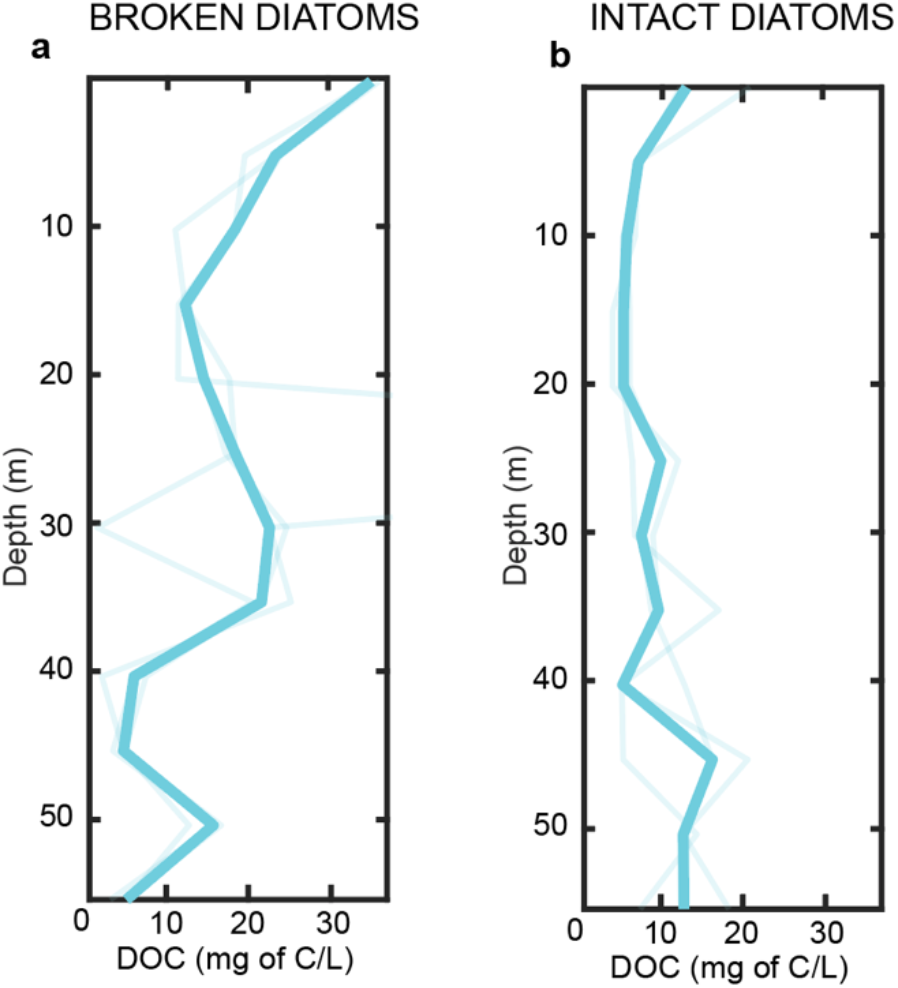
DOC depth profiles of particles seeded with broken diatoms and intact diatoms. **a)** DOC depth profiles of particles seeded with broken diatoms. The thicker line is the mean of three replicate devices each containing n =4 biological replicates, 12 particles in total. **b)** same as a, but of particles seeded with intact diatoms.

### A highly localized oxygen landscape driven by divergent carbon reserves

Oxygen is perhaps one of the most dynamic state variables across a marine snow particle actively being colonized by heterotrophs. As such bacteria consume the oxygen around them, they begin to self-limit their own aerobic growth. At the scales of the microorganisms themselves, this can happen across a particle or across even a microcolony itself (Extended Data Fig. 5–8) For facultative anaerobes, this highly localized oxygen landscape translates to a locally heterogeneous metabolic population and increases in niche functions ^35,36^. Indeed, the evolution of a particle’s oxygen landscape is apparent in these experiments (Extended Data Fig. 5, 7). At the early time points (analog to depth in the real ocean) of the experiments, the vast majority of colonies are well-oxygenated at concentrations close to the saturation conditions of the bulk seawater flowing around the particle. As the particle descends, however, these colonies become progressively more suboxic (∼20 m; Extended Data Fig. 5, 7). However, this decrease in oxygen concentration causes the organisms themselves to slow down via their regulatory network, which in turn permits the oxygen concentrations to slightly increase before continuing to decrease. This stage coincides with very little change of oxygen across a particle or microbial growth (Extended Data Fig. 5–8). At the end of the progression, the colonies across the particle are close to fully anoxic, with the entire microbial metabolism supported solely by the leaking carbon from diatom cells. Interestingly, both the colonies in close proximity to the particle edge and those nearer to the center can become anoxic in this system, even if those further interior can more readily experience anoxia due to the greater distance to resupply oxygen diffusively from the bulk seawater. Notably, the oxygen concentrations observed by the colonies themselves are nearly indistinguishable from the concentrations at the equivalent particle radius where no colonies develop, even though the colonies themselves are the sinks of oxygen (Extended Data Fig. 5–8). This occurs because at the very small spatial scales, the colony’s oxygen equilibrates quickly (less than a second) with its immediate surroundings via diffusion.

**Fig. 4.**
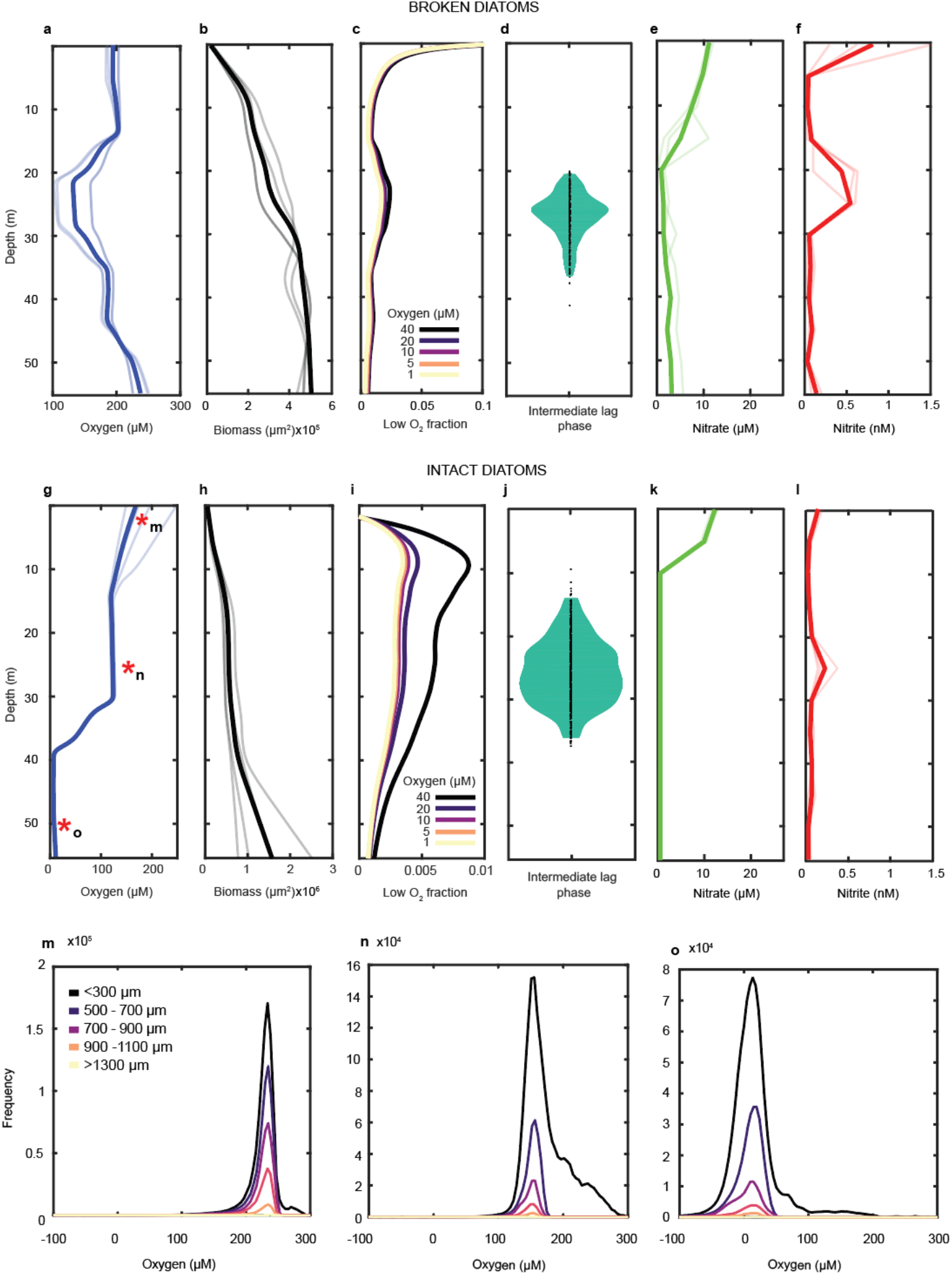
Oxygen, biomass accumulation and nitrate and nitrite depth profiles. For particles seeded with broken diatoms: **a)** Oxygen change at the colony level due to colony respiration. The thick line is the mean value of four particle replicates; each indicated with a lighter color. **b)** Biomass accumulation as determined by colony cross-sectional area. **c)** Fraction of total colony area that experiences different oxygen conditions. Each line shows the area where oxygen is less than the shown value. **d)** Intermediate lag phase distribution. **e)** Nitrate evolution in the bulk seawater. **f)** Nitrite evolution in the bulk seawater. For particles seeded with intact diatoms, panels **g–l** correspond to the broken diatom counterparts. **m–o)** Histograms of oxygen concentrations within colonies at 5, 20, and 40 m, respectively. The colonies are distinguished by their locations in the particles. Asterisks in **g** denote the time points of these panels.

**Fig. 5.**
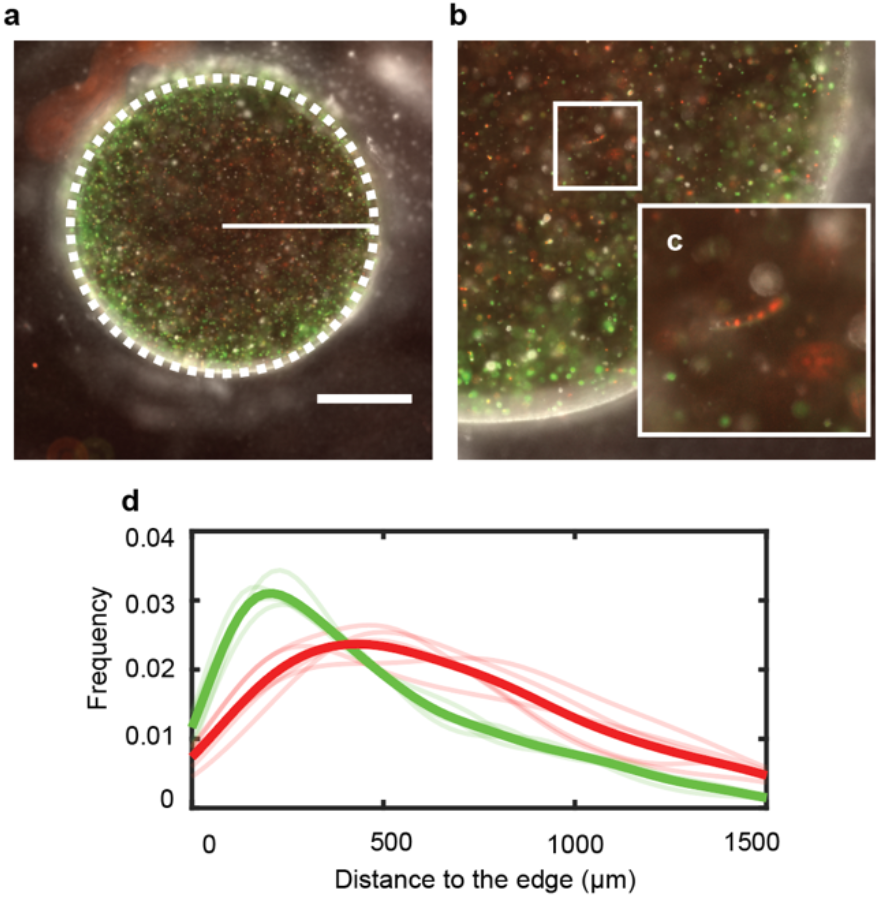
Spatial expression of NarK and NirS genes and diatoms. **a)** Particle seeded with *Pseudomonas aeruginosa* PAO1 NarK-GFP and PAO1 NirS-dsRed and *Chaetoceros affinis*, the dashed white line identifies the particle edge while the solid line indicate the radius, indicated the radial sampling of the gene expression within the particle. **b)** Close up of quarter of the particle showing the diatoms and denitrifying colonies **c)** Inset shows in detail the intact diatom within the particle with the chlorophyll fluorescing naturally. **d)** Probability density functions of the expression of NarK-GFP and NirS-dsRed shown for each replicate, with an average overlain with a thicker curve. The white scale bar in **a** denotes 1000 μm.

In particles seeded with broken diatoms approximately one mode of marine snow generation by zooplankton repackaging, the microcolony respiration causes oxygen to reach its lowest concentration at 20 — 30 m effective depth (Fig. 4 a) before oxygen increases again as the biomass enters stationary phase (Fig.4 b). This stage coincides with a short plateau of relatively constant average oxygen of ∼100 μM (Fig. 4 a) across the colonies. This intermediate plateau in oxygen consumption corresponds to a temporary increase of local biomass experiencing (Fig. 4 c) hypoxic (<40 μM), microoxic (<10 μM), suboxic (<5 μM) and nanoxic (<1 μM) conditions ^37^. That is, even though the median colony does not change in its observed oxygen concentration, the number of colonies (although still a vast minority of cells) experiencing localized oxygen low enough to permit denitrification does increase. For a facultative heterotroph, the decrease of oxygen substantially impacts its growth rate and metabolic machinery. To pinpoint when this change in ‘metabolic gear’ occurred, we analyzed the growth phases of all the colonies across a particle to isolate the moment in which an intermediate lag phase emerges in response to oxygen drawdown (Fig. 4 d). This intermediate stage of the colony growth, while relatively short occurring over 9.6 ± 4.1 m of sinking (mean ± stdev) denotes the point when the local metabolic machinery is being rebuilt to adjust to local oxygen availability. This intermediate plateau of the growth notably has an impact in the water column, giving rise to a simultaneous visible nitrite peak at the same depth. When finally the bacteria reach late stationary phase, having exhausted all labile carbon resources at >30 m depth (Fig. 4, c), oxygen penetrates again within the particles and each colony equilibrates with oxygen in bulk seawater. Notably, there are many hypoxic and lower colonies initially in this experiment, corresponding to the greatest rate of carbon leakage from the broken diatoms (and the highest nitrite concentrations measured).

Particles mimicking intact diatom aggregates, however, display a similar yet importantly different metabolic evolution. After an initial decrease of oxygen driven by the initial exponential growth of the biomass accumulation (Fig. 4 g,h), colonies subsisting under local microaerobic conditions (<40 μM) expand much more than the biomass that experience other low oxygen conditions (Fig. 4 i). After an initial exponential growth phase, the bacteria slow and enter an intermediate lag phase contemporaneous with the intermediate intraparticle oxygen plateau. Like the particles seeded with broken diatoms, this intermediate lag phase of the biomass growth occurs when the colonies reach a local oxygen average concentration around 100 μM. These cells also display a wider lag phase distribution (12.7 ± 5.0 m mean ± stdev) compared to those in association with broken diatoms. At the end of this intermediate lag phase, the colonies restart their metabolism and enter a second exponential growth phase, now nearly fully anoxic, and still supported by the available carbon continuously provided by the extant diatom cells. Like the particles seeded with broken diatoms, these too indicate that the metabolic activities are such that nitrite only accumulates in the bulk at the peak of the of the lag phase distribution. The particles seeded with broken diatoms show a significantly shorter lag phase compared to the particles seeded with intact diatoms (two-sample t-test, p = 1.3 × 10^−22^).

When comparing the experiments conducted with intact diatoms and their broken counterparts, general trends emerge with some notable distinctions. Under both scenarios, multiple microbial growth phases appear that cause oxygen to be reduced at the scale of a colony. The same opposing forces at play in an ODZ — biological consumption of oxygen and physical resupply — yield a time (ergo, depth) at which oxygen is a minimum. Oxygen is consumed more rapidly initially with the broken cells leaking much of their organic contents as a pulse, stimulating growth but leaking DOC into the surrounding seawater. An initial leak of nitrite is also observed here in all broken diatom experiments (Fig. 4 f). However, colony growth stagnates quickly as the organic matter contained within the phytoplankton is not retained in the particle. In contrast, intact cells permit a slow leakage of organic material that delays colonization but ultimately creates an environment conducive for a greater microbial carrying capacity and thus higher oxygen drawdown and rates of denitrification. In fact, both broken and intact diatoms yield similar progressions for the first half of the experiments, but in the broken trials, labile organic matter is exhausted earlier whereas in the intact ones, the bacteria at the edge have sufficient resources of both reductant and oxidant to continue to colonize and thrive. The only cells growing at the latest time points in the intact experiments are those at the edge, but they are growing aerobically given the contraction of the suboxic area and constant size of the anoxic zone.

### Spatial segregation of nitrate and nitrite reduction

To understand more mechanistically how denitrification in marine snow manifests, the impacts for water column chemistry, and link gene expression with location across the particle and highly localized oxygen conditions, we additionally mixed diatom cultures with denitrification-reporting strains of the ubiquitous bacterium and facultative denitrifier *Pseudomonas aeruginosa*, an organism with the full complement of enzymes to reduce nitrate to dinitrogen. Whereas this organism is found in the ocean itself ^38–40^ but is typically studied as an opportunistic human pathogen, here it is used as a representative model for similar marine denitrifiers, many of which are also gamma proteobacteria (e.g., *Marinobacter, Alteromonas*, and *Idiomarina* spp.) ^41,42^.

The different steps of denitrification are each subject to different regulation by oxygen and other stimuli. Typically, nitrate reduction, the first step of canonical denitrification, can manifest at a higher oxygen tolerance than later steps, whereas the ultimate step of nitrous oxide reduction, is the most severely inhibited by oxygen ^1,43^. Under this framework, the spatial segregation of the denitrification steps across a particle can be anticipated because higher oxygen concentrations will be maintained at the periphery. Within our particle experiments harnessing *P. aeruginosa* as a laboratory model of real world marine denitrifiers, the diatoms support sufficient growth of the bacterial colonies and the commensurate oxygen drawdown that denitrification activates as measured by fluorescent reporter constructs for the promoters of the nitrate or nitrite reductase enzymes (Fig. 4). Notably, the same phenomenon manifests whether growth is supported by the large chain-forming diatom *C. affinis* or by the smaller pennate diatom *P. tricornutum* (Extended Data Fig. 9).

In both experiments with different diatom sources, active denitrification (both nitrate and nitrite reduction) is observed across the particle, and moreover, in both, nitrate reductase regulation is skewed toward the particle edge whereas nitrite reductase reaches its peak regulation inward of the edge. We explain this as arising from the dual control of both higher oxygen and nitrate toward the periphery, activating nitrate reductase instead of nitrite reductase. This spatial differentiation of activity between these two steps can yield a substantial impact on the bulk, as the nitrite product of the first step diffuses equally inward and outward. Whereas later denitrification steps can reduce the nitrite in the particle center, whatever diffuses outward can leak into the bulk environment and modify the water column chemistry and subsequent planktonic biological activity in response.

## DISCUSSION

### Implications for the fixed nitrogen budget

The marine fixed nitrogen budget has been debated for decades, whether it is balanced or not ^44^, and what the climactic implications are for a global world if not. Additional pathways and mechanisms have been revealed, but the general budgets maintain fixed nitrogen loss by denitrification (and anammox) occurs exclusively within sediments and in the ocean’s oxygen deficient zones. Here we have shown particles found throughout the water column can be another marine environment that can readily support denitrification and other anaerobic metabolisms. The expansion of denitrification capabilities on particles can dramatically increase the gross fixed nitrogen loss rates in the global oceans ^7^. Whether the net loss rates, further suggesting a pronounced imbalance of sources and sinks, are in actuality greater than current estimates or are counterbalanced by additional diazotrophy in the oceans’ interior, remains to be evaluated. Yet, increased denitrification in the interior driven by particles can also explain the abundance of heterotrophic nitrogen fixers within particles and at depth ^45,46^. Importantly, given that particles themselves are small and widely distributed, a dramatic nitrate deficit compared to phosphate indicative of denitrification, akin to the ODZs, is not to be expected. Interestingly given the experiments with intact and broken diatoms, the composition of a particle may not be strictly important for denitrification to occur as long as there remains some labile organic carbon. Nonetheless, to fully resolve the importance of marine particles for the fixed nitrogen budget, their complete size spectrum must be fully investigated for denitrification feasibility.

Nitrate and nitrite in such a system may also not be anticipated to be fully reduced to dinitrogen, but rather the denitrification cascade could be halted at nitrous oxide (N_2_O) given the severe inhibition of oxygen for nitrous oxide reductase. Moreover, in the real ocean, a particle is likely seeded by a diverse assemblage of organisms consisting not only of facultative denitrifiers but also autotrophic nitrifiers benefitting from the ammonification of the organic matter released from the phytoplankton. Aerobic nitrifiers too have been found to be associated with marine particles ^47^, and anammox has been shown to rely on the ammonium released from organic matter degradation ^48^. Particles would easily permit the low-oxygen nitrification pathway that dramatically increases N_2_O yields ^49,50^ to activate. Indeed, consideration of particles could also resolve an additional mystery related to the difference between N_2_O yields as implied from global model inversions ^51^ vis-à-vis measured in situ in the ocean and in cultures ^52,53^. Marine samples seemingly show lower N_2_O yields at a given oxygen concentration than inverse models, but if the metabolism were taking place in microsites with different oxygen conditions than the bulk, the disparate observations could be reconciled.

### On the formation of the primary nitrite maximum

In all experimental conditions — intact or damaged diatoms paired with marine bacterial isolates, the natural xenic phycosphere, or the laboratory model *Pseudomonas* — a transient nitrite accumulation is observed in time. Because here time is an analog for depth, we can transpose the axis, the result of which is a layer of nitrite buildup that corresponds to ubiquitous primary nitrite maximum (PNM) across the world’s oceans. Whereas the measured increase in nitrite here is small (approx. a nanomole per liter), this arises from just 4 biologically viable particles in the nearly 15 mL that flows past them per daily measurement. The actual particle concentrations in the ocean are orders of magnitude more abundant, increasing the impact intraparticle nitrate and nitrite reduction by denitrifiers have on water column chemistry. The PNM has classically been considered to arise from light limited phytoplankton incapable of fully reducing nitrate to ammonium for assimilation ^54^ and/or inefficient coupling of ammonium and nitrite oxidizers during light-inhibited nitrification ^55^ This work notably does not exclude these leading hypotheses behind the formation of the PNM as each of these causes would still yield nitrite accumulation in a similar layer. Yet, whereas these two previous hypotheses have been discussed at length, we focus here on the potential role of particle-based denitrification in contributing to the PNM.

The secondary nitrite maximum manifests from denitrification in the anoxic oxygen deficient zones of the eastern tropical Pacific and Arabian Sea wherein nitrate reduction outpaces nitrite consumption ^56,57^. Here we suggest an analogous phenomenon in shallower subsurface waters in contributing to the ubiquitous PNM. A transect across the eastern tropical North Pacific reveals the PNM aligns with the highest particle concentrations (as inferred by increased beam attenuation) (Fig. 6). Moreover, stations where the particle concentrations are higher manifest more nitrite accumulation in this PNM layer (Fig. 5b). In terms of quantity, size, and total biovolume, the steep density gradient (pycnocline) retards sinking rates and traps particulate material ^58–60^. These particles, derived from the surface, act to support increased heterotrophy in the shallow subsurface. Indeed, the depths of the steepest pycnocline align identically with the PNM (Fig. 6). Yet, the same conditions that trap particles at these depths would increase the rates of nitrification as well. Further, because of the deep chlorophyll maximum just above this layer, the contribution of light-limited phytoplankton harnessing insufficient photons to efficiently reduce nitrate to ammonium via nitrite cannot be excluded. Yet, none of these pathways to produce nitrite at the PNM needs be mutually exclusive. Rather, a combined effect from all is the most likely reason due to the intrinsic alignment of the pycnocline with the deep chlorophyll maximum, i.e., the DCM naturally arises just below the mixed layer where cells have less light but greater access to nitrate. Given all these overlapping phenomena, we propose that particle-based denitrification complements leaky phytoplankton and decoupled nitrification as a pathway leading to the rise of the primary nitrite maximum (Fig. 5c).

**Fig. 6.**
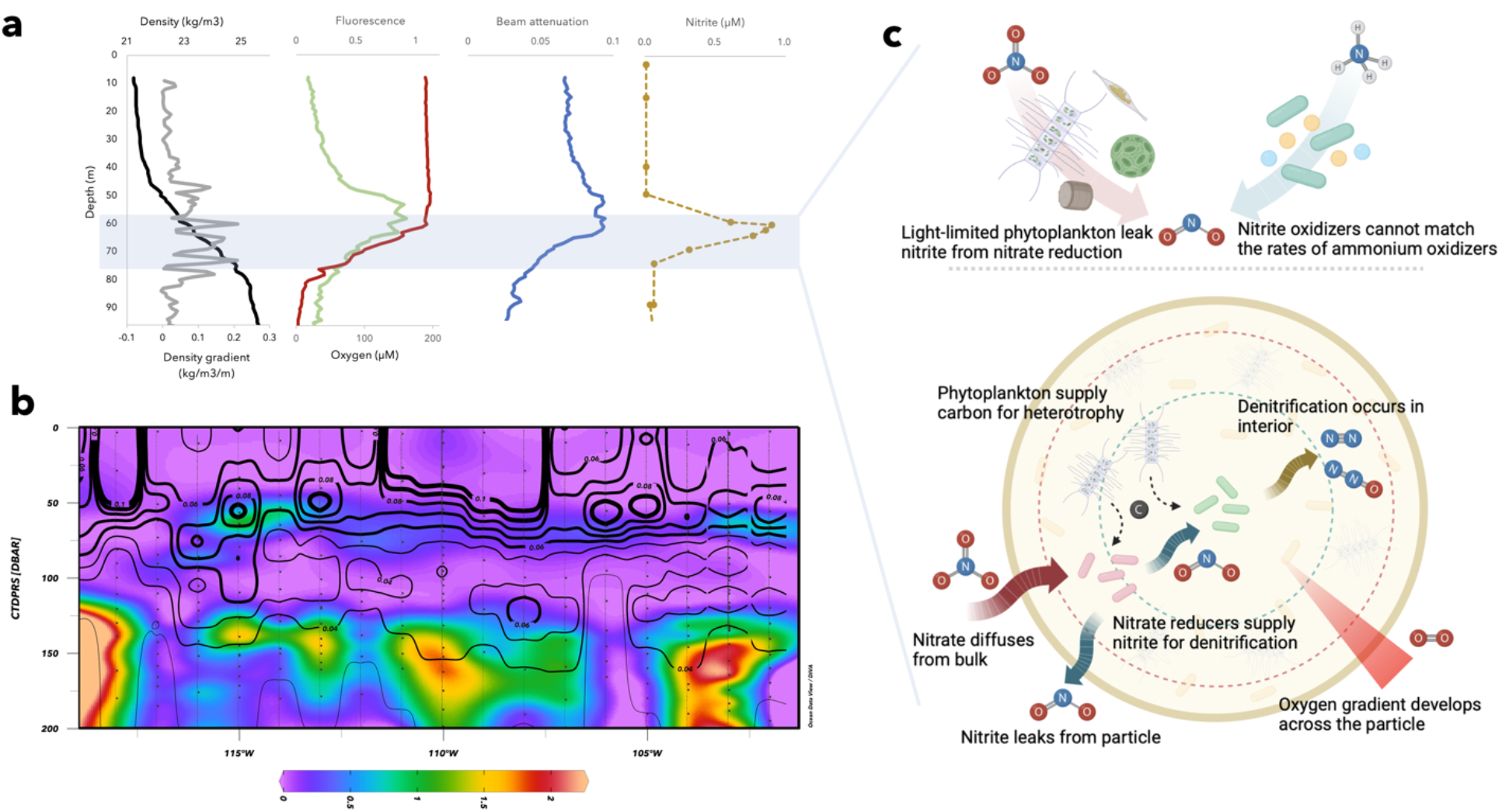
Particles and the primary nitrite maximum. **a)** profiles of density (black), vertical density gradient (grey), fluorescence (green), oxygen (red), beam attenuation (blue), and nitrite (brown). The profiles show that the depths of accumulation at the primary nitrite maximum (noted in the blue box) align with the depths of maximal density gradient. Additionally, the primary nitrite maximum is just deeper than the fluorescence maximum of chlorophyll and aligns with maximal beam attenuation. **b)** zonal section of nitrite and beam attenuation from the R/V *Falkor* FK180624 cruise showing nitrite (color) and overlain by beam attenuation (black contours). **c)** potential pathways that lead to the production of nitrite at the primary nitrite maximum. The top two are well-established, light limited phytoplankton reducing nitrate inefficiently and ammonium oxidizers operating faster than nitrite oxidizers in nitrification. The bottom schematic relays what we here assert as plausible, i.e., that denitrification within particles composed of phytoplankton detritus can be separated in space due to gradients in oxygen across a particle. Nitrate reduction occurs at the edge, permitting nitrite to diffuse inward where it may be further denitrified to N_2_O and N_2_ and outward where it may be released to the water column.

### Broad implications

Taken together, the experiments combining different diatoms and bacteria highlight how heterotrophic colonies grow and denitrification readily emerges within a marine snow particle supported solely by the organic lysate leaking from phytoplankton. Impressively, even the broken diatoms are not immediately and completely solubilized; rather, these particles similar to what may be repackaged by zooplankton retain organic carbon far longer than the approx. 20-minute time scale to flush the particle via diffusion. Intact diatoms retain their carbon even longer, permitting more growth, greater draw down of oxygen, and correspondingly higher denitrification potential across over 10 days of sinking. The elucidation of this phenomenon coupled to the prevalence of marine particles globally and throughout the water column, can have far ranging implications for bulk ocean chemistry, the fixed nitrogen budget, and potentially nitrous oxide production. How these large budgets are resolved to include particle denitrification, and potentially other metabolisms favored in low-oxygen environments like diazotrophy, sulfate reduction, and methanogenesis, remains to be determined with directed and specific efforts ^7^.

The dynamic micro-suboxic and micro-anoxic sites that develop across a particle during its sinking can further help reconcile additional outstanding questions like the dominance of the nitrite oxidoreductase (NXR) protein in the oceans ^61^. Given how nitrite leaks from each of these particles, it must either accumulate or be consumed. In the oxygenated bulk ocean, aerobic nitrification is the most likely pathway, and one catalyzed by NXR. Notably, this mechanism also implies a recycling between the two oxidized forms of fixed nitrogen, nitrate and nitrite, rather than the source of nitrite being ammonium from organic matter remineralization. As such, this mechanism would permit a decoupling of the nitrification steps. Whereas the magnitude of the impact of denitrification in sinking marine snow to the PNM remains to be fully resolved, from our experimental particle and field data, denitrification cannot be excluded, at least in regions of higher productivity, as contributing to nitrite accumulation at the PNM.

## METHODS

### Isolation and phenotypic assay

Five nitrate-reducing heterotrophs — *Alterosomas* sp.; *Phaeobacter* sp.; *Idiomarina* sp.; *Virgibacillus* sp.; *Marinobacter* sp. — were isolated from surface seawater collected in the Sargasso Sea (collected and filtered in June 2014 at 40°67.4269’ N, 70°53.3153’, depth of 60.5 m, with salinity of 33.16 and temperature of 11.7ºC), and from the phycosphere of our *Chaetoceros affinis* culture. First the seawater was enriched with filter sterilized (0.22 μm) diatom exudates for five days at room temperature (21°C). 5 mL of diatom culture was centrifuged at 7000 rpm for 4 minutes at room temperature and resuspended in 500 μL of sterile seawater to concentrate the diatoms in a small volume. 100 μL of the enriched seawater and 100 μL of the diatom-concentrated seawater were then inoculated on agar plates (1% agar – VWR Agar for bacteriology) supplemented with Marine Broth 2216 (VWR) (MB) and incubated at room temperature (21°C) for 5 days under aerobic conditions. Single colonies were collected and identified using 16S rRNA amplicon sequencing.

Each colony was resuspended in 5 mL of MB amended with 10 mM of nitrate, incubated for 48 h at room temperature under anaerobic conditions in a glove box (Coy Laboratory Products, Grass Lake, USA) with defined atmosphere, nitrogen (N_2_):hydrogen (H_2_) atmosphere (97:3). To determine their denitrification capability, first nitrite was measured with the standard colorimetric Griess reaction 62. Second, we recorded nitrous oxide emissions (Extended Data Fig. 1) to confirm the complete denitrification capability of the community. We seeded agarose particles with the marine isolates (final OD = 0.0002) and incubated them under continuous flow of aerobically sparged (compressed air 10 mL/min, volumetric flow of 0.01 mL/min) MB in an air-tight millifluidic device. A nitrous oxide microsensor (Unisense, Aarhus, Denmark) with steel needle was inserted in the millifluidic outlet to record the dynamic production and consumption of nitrous oxide and confirm a complete denitrification pathway (Extended Data Fig. 1).

### 16s rRNA identification

Nineteen colonies isolated were resuspended in sterile PBS solution and the DNA was extracted using the DNeasy Powerlyzer Microbial Kit protocol (Qiagen, Hilden, Germany). After DNA extraction, the purity of the DNA was quantified with NanoDrop UV/Vis spectrophotometer (NanoDrop Technologies, Wilmington, DE). The DNA was then amplified with Quick-Load Taq 2X Master Mix (New England BioLabs, Ipswich, MA) and universal bacterial 8 F/1492 R primers. The samples were sent to Eton Bioscience for Sanger DNA sequencing using primers 8 F and 907 R. After the 16S rRNA identification and the phenotypic essay, 5 unique isolates at the genus level from the 19 candidates in total were obtained to use in the further experiments.

### Inocula preparation

*Chaetoceros affinis* and *Phaeodactylum tricornutum* were purchased from NCMA (Bigelow, National Center of Marine Algae and Microbiota, East Booth Bay, Maine). Once the diatom cultures arrived, they were immediately transferred to L1 media ^63–66^. Prior to seeding the diatoms in the agarose, the diatom culture was grown for 5 days, *C. affinis* at 26°C and *P. tricornutum* at 19°C. The *C. affinis* intact diatoms were prepared in the following way. When the diatoms reached an OD of 1.5 at 320 nm, 5 mL of the culture was then filtered using a Falcon® 40 μm Cell Strainer to collect intact cells and resuspended in 1 mL of sterile seawater. 250 μL of *C. affinis* intact diatom suspension were finally inoculated in 1 mL of low melting agarose and gently vortexed to uniformly distribute them in the hydrogel. To obtain the broken diatom, 1 mL of diatom culture was sonicated for 1 minute, power set at 60% (Branson Ultrasonics™ Microtips for 200 Watt Sonifier; Branson Ultrasonics™ Sonifier™ SFX250/SFX550 Cell Disruptors). 250 μL of *C. affinis* broken diatom suspension were finally inoculated in 1 mL of low melting agarose and gently vortexed to uniformly distribute them in the hydrogel. The *P. tricornutum* diatoms were filtered using a Falcon® 40 μm Cell Strainer to collect intact cells and resuspended in 1 mL of sterile seawater. 250 μL of *P. tricornutum* suspension were finally inoculated in 1 mL of low melting agarose and gently vortexed to uniformly distribute the cells in the hydrogel.

Five bacterial species — *Alterosomas* sp.; *Marinobacter* sp.; *Phaeobacter* sp.; *Idiomarina* sp.; *Virgibacillus* sp. — were incubated overnight at 30°C in 5 mL of Marine Broth 2216 (VWR) in a shaking incubator at 220 rpm. 200 μL of each overnight culture was resuspended in 20 mL of Marine Broth 2216 (VWR) in an Erlenmeyer flask in shaking incubator at 220 rpm for 3 h until each species reached an optical density of 0.15. 12 μL of each culture was inoculated into 1000 μL of low gelling agarose containing 10 μL of PyroScience oxygen nanoprobes (Oxnano; 10 mg/mL) and diatoms.

*Pseudomonas aeruginosa* PAO1 NarK fusion reporter and *P. aeruginosa* PAO1 NirS fusion reporter ^35^ were incubated in a shaking incubator at 220 rpm at 37°C overnight. 200 μL of the overnight culture, was then resuspended in 20 mL of sterile LB media to reach 0.15 OD after 2 h of incubation in a shaking incubator at 220 rpm and 37°C. 12 μL of each reporter strain was resuspended in low melting agarose containing 10 μL of Oxnano (10 mg/mL) and diatoms with the same density as the marine isolates experiments described above.

### Millifluidic experiments

Once the low melting agarose was seeded with oxygen sensitive nanoparticles, bacteria, and diatoms, it was cast into 1.5 mm radius disks 700 μm in height and prepared as described previously ^35^. The hydrogel particles were then enclosed in a millifluidic device as described before ^35^, each containing 4 live seeded particles and one inert one upstream for calibration purposes. All the sinking particle experiments were performed with the same seawater, which was collected in October 2019 at 39°46.406’ N, 70°53.065’ W, bottom depth of 1578 m, with a salinity of 35.67, sea surface temperature of 23.3°C, and below detection limit chlorophyll. The seawater was filter sterilized with a 0.20 μm vacuum filtration cup (VWR, 10040-468). In order to confirm the absence of any nitrate or nitrite, prior to performing the experiments, it was first tested for the presence of NOx^−^ species with the chemiluminescence method described below. The seawater was then amended with concentrated stocks of sodium phosphate monobasic (S3522-500G, Sigma Aldrich) to obtain a final concentration of 1 μM and sodium nitrate (SS0680-500GR VWR) to obtain a final concentration of 10 μM.

The seawater was then transferred to acid-washed and autoclaved 125 mL serum bottles and sparged continuously with air (10 mL/min). The volumetric flow through the millifluidic devices was maintained at a rate of 0.01 mL/min using a syringe pump (Harvard Apparatus PHD Ultra Syringe Pump), with 15 mL syringes (BD Luer-Lok tip) with in-line filter to sterilize the sample (Acrodisc® Syringe Filters with Supor® Membrane, Sterile - 0.2 μm, 25 mm). The seawater was sampled and stored in 40 mL VOA (ThermoFisher Scientific Catalog # 40-EPAVCSA). Each vial was washed with HPLC grade acetone (34850-1L, Sigma Aldrich) and then combusted in an oven at 500°C for 8 h. Milllifluidic devices and inlet and outlet tubes (Viton®, fluoroelastomer, High-Temperature Soft Rubber Tubing, Mc Master-Carr) were tested for the release of background dissolved carbon, and no detectable dissolved carbon was confirmed. Millifluidic devices with empty agarose particles were also tested for the release of background dissolved carbon, and no detectable dissolved carbon was found. Experiments were conducted in the dark, simulating the export of cells below the euphotic zone when they can no longer photosynthesize.

### Elemental carbon analysis

To monitor the change in total dissolved organic carbon (DOC), we used an Elementar Vario-EL analyzer modified to introduce liquid samples. We constructed our DOC calibration curve using a potassium hydrogen phthalate standard solution (LabCem, USA). Samples were analyzed within 5 hours of collection. The samples were first filtered through 0.22 μm glass fiber filters (Kinesis KX, Canada) and diluted 5 times with in-lab produced milli-Q water. The samples were then acidified with 3 drops of 37% hydrochloric acid (Acros Organics, USA) and analyzed. The analysis program included triplicate flush and injections with 0.2 mL per injection. After each run, a flush sequence was conducted to eliminate cross-contamination between samples.

To assess the carbon content in the initial diatoms used, we measured dry mass organic carbon using an Elementar Vario-EL analyzer modified to introduce solid sample equipped with non-dispersive infrared detector (NDIR). We constructed our calibration curve using potassium hydrogen phthalate salt (Elementar, Germany). A small diatom sample was first dispersed in 250 μL of seawater. Then, the samples were transferred into clean tin capsules (Elementar, Germany) to prepare for total solid organic carbon (SOC) analysis. Clean empty capsules were first weighed using microbalance (Mettler Toledo, USA), then the diatoms were transferred. The capsules were then allowed to dry in a 40°C oven for 24 hours. The samples were weighed again, and the difference was computed. A blank was prepared to account for seawater matrix. Last, the capsules were acidified with two drops of 12% hydrochloric acid (Acros Organics, USA) to liberate any inorganic carbon and analyzed. The total collected dry mass for broken and intact diatoms were 0.385 and 0.328 mg, respectively. On a per-particle basis, the broken diatom particles were seeded with 0.0585 mg of total carbon whereas the intact ones were seeded with 0.0497 mg of C per particle.

### Nitrate and Nitrite measurements

The nitrate was measured using a standard chemiluminescence approach by quantifying the total NO_x_^−^ species via chemical reduction to NO, performed with hot acidified vanadium (III). Once the NO_x_^−^ is completely transformed to NO and quantified with a Teledyne T200 NOx analyzer ^67,68^. Nitrite was quantified using 96 well plates with the Griess colorimetric assay ^62^ and a plate reader (UV-Vis, Tecan, Spark) measuring the absorbance at 543 nm. 10 μL of combined Griess reagent (sulfanilamide and N-napthylethylenediamine) was added to 200 μL of seawater using the automatic injector module, incubated for 1 h at room temperature in the dark, and measured performing a lambda scan from 400 to 700 nm to obtain the full spectrum of absorbance for a more precise readout at sub-nanomolar concentrations (Extended Data Fig. 2).

### Image acquisition and analysis

All the images were acquired with a Nikon Eclipse Ti-2 microscope with an Andor Zyla 4.2 sCMOS VSC-06626 camera, 10x objective (Plan fluor), bright field and CY5 filter cube set (for the oxgen nanoprobe fluorescence), FITC for GFP and CY3 for dsRED. A time series of images at an interval of 1 h was continuously acquired until colonies stopped developing, for a total of at least 13 days. At each time point, a bright field image with a pseudo dark field setting was acquired together with the fluorescence image (excitation: 590–650 nm, emission: 662–737 nm) for the oxygen sensitive nanoparticles. CY3: excitation: 532 to 557 nm, emission 570 to 640 nm while FITC excitation 465 to 495 nm and emission from 525 to 515 nm. These two were only used on a separated experiment for the detection of the fluorescence reporters of Nar and Nir genes, respectively GFP and dsRed fluorescence proteins. All images were analyzed using MATLAB (R2016b) using the image analysis toolbox. At the end of each experiment, the particles were exposed to chloramphenicol (35 to 50 μg/mL) to interrupt the metabolic activities of the bacteria. Seawater amended with chloramphenicol and sparged with N_2_ was flushed within the millifluidic device to obtain the anaerobic calibration of each particle. Multiple images were acquired, and using the NIS-Elements software, a profile of nanoparticle fluorescence across the particles was plotted during the image acquisition. When the fluorescence profile showed complete saturation of the nanosensor fluorescence (indicating full anoxia of the entire particle), these images were selected for the anaerobic calibration endmember. The same process was repeated, but with air-sparged media, to obtain the air-equilibrated particle calibration endmember. Notably, each particle has its own set of aerobic and anerobic images for calibration of the nanosensor to account for any heterogeneity in the nanoparticle distribution. Later, during the image analysis script, the calibration particles were divided into spatial zones as a function of radius to obtain a median value of the fluorescence nanosensor signal and create an optimal highly resolved local calibration (Fig. 1 i, and Extended Data Fig. 3).

Using the elongation shape factor, corresponding to the square root of the ratio of the two second moments of an object around its principal axes ^69^, the diatoms were segmented and excluded from the fluorescence image in order to exclude their potential interference (by chlorophyll) with the oxygen nanoparticle signal (Extended Data Fig. 4 a-d). A custom backtracking algorithm enabled us to follow each individual colony expansion while simultaneously quantifying the oxygen concentration change across the particle and within each individual microcolony (Extended Data Fig. 4 e). The edge of the hydrogel particles was detected using the first image (Extended Data Fig. 4 e). The images were enhanced using *imadjust* function, then using a Gaussian filter with a smoothing kernel with standard deviation of sigma = 20 followed by a global threshold and connected component analysis to fill the entire particle area (Extended Data Fig. 4 e). Finally using the built-in function *regionprops*, the edge of the particle was identified. Using the built-in *edge* function followed by Otzu binarization. The binarized image is first dilated and then filled using a flood-fill operation, allowing bacterial colonies to be identified. For each individual colony, its position and distance from the particle edge was recorded (Extended Data Fig. 4 e). For every colony at each time point, its cross-sectional area and median oxygen value (obtained using the local calibration values) was stored. The distribution of the signal of the oxygen nanosensor of each colony and of the particle excluding the colonies was stored as equivalent oxygen values (obtained using the local calibration values) at each time point (Fig. 1, Extended Data Fig. 3).

