## SupplementaryInformation for "Sinking diatom aggregates provide carbon to drive microscale denitrification in a bulk oxygenated ocean"

**Table of content:**

1. Extended Data Fig. 1 Nitrous oxide quantification
2. Extended Data Fig. 2 Nitrite quantification with Griess method.
3. Extended Data Fig. 3 Aerobic and anaerobic calibration of in situ oxygen nanoprobes in the particles.
4. Extended Data Fig. 4 Step-by-step image analysis algorithm.
5. Extended Data Fig. 5 Probability density function of oxygen change at colony level of particles seeded with broken diatoms.
6. Extended Data Fig. 6 Probability density function of oxygen change at particle level of particles seeded with broken diatoms.
7. Extended Data Fig. 7 Probability density function of oxygen change at colony and at the particle level of particles seeded with intact diatoms.
8. Extended Data Fig. 8 Probability density function of oxygen change at colony and at the particle level of particles seeded with intact diatoms.
9. Extended Data Fig. 9 Spatial expression of NarK and NirS genes for experiment with *Phaeodactylum tricornutum*.
10. Extended Data Fig. 10 Nitrite evolution in the water column of particles seeded with *Pseudomonas aeruginosa* and *Phaeodactylum tricornutum* or *Chaetoceros affinis*.
11. Extended Data Fig. 11 Nitrate evolution in the water column.
12. Extended Data Fig. 12 Nitrite evolution in the water column.
13. Extended Data Fig. 13 DOC evolution in the water column.

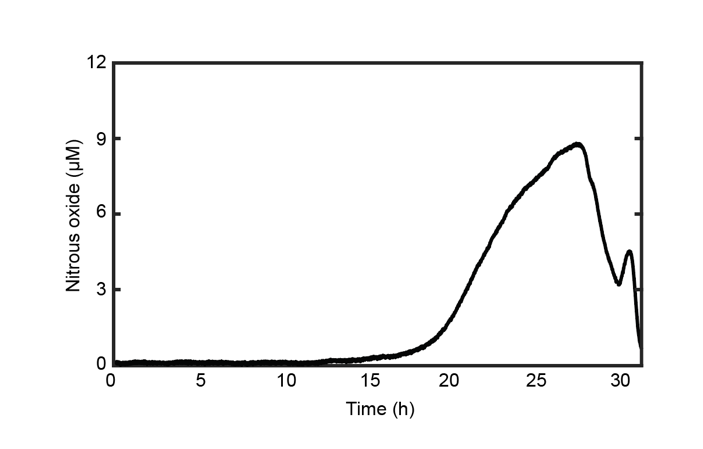

**Extended Data Fig. 1 Nitrous oxide quantification.** Nitrous oxide production and consumption of four particles seeded with the marine isolates used in this study, confirming denitrification capability. The signal was measured with a microsensor inserted at the outlet of the millifluidic device. The media used for this test was Marine Broth 2216 amended with 400 µM nitrate.

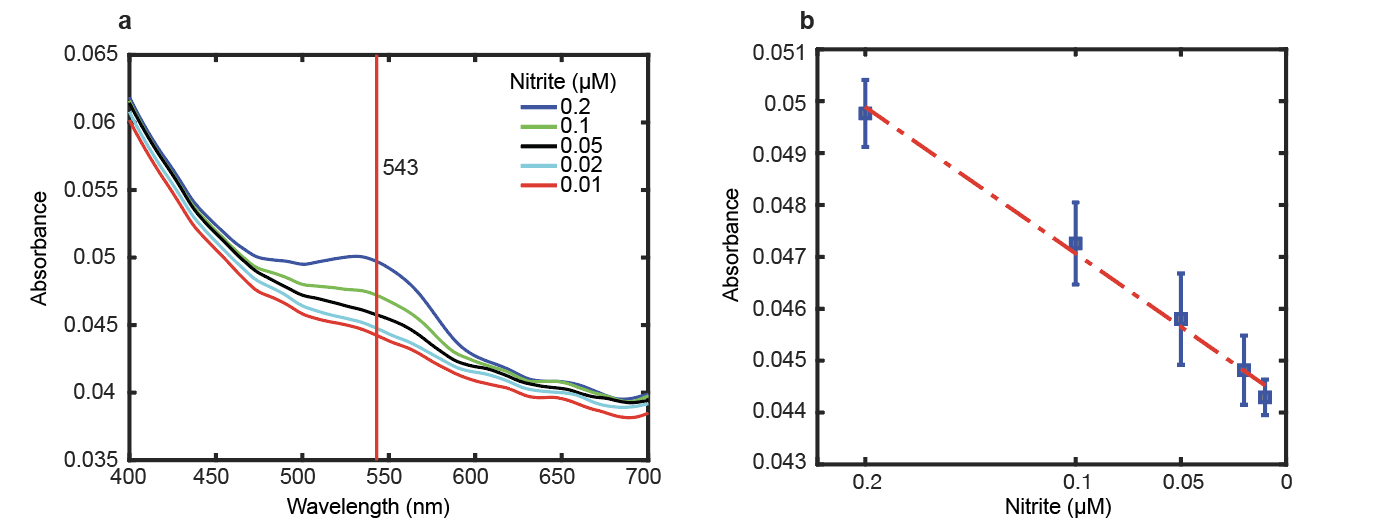

**Extended Data Fig. 2 Nitrite quantification with Griess method. a)** Mean signal of five technical replicates of lambda scan 400–700 nm. The red line indicates the 543 peak, corresponding to the maximum absorbance of Griess test solution. **b)** The squares represent mean value of five technical replicates of absorbance peaks; the error bars represent the standard deviation. The linear best fit equation is shown.

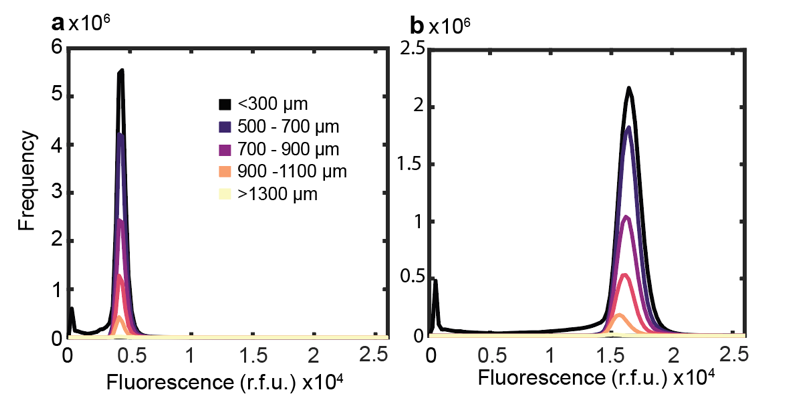

**Extended Data Fig. 3 Aerobic and anaerobic calibration of in situ oxygen nanoprobes in the particles.** A representative particle is shown for **a)** aerobic and **b)** anaerobic calibration. Each line is color coded based on a defined zone as a function of spatial location in the particle. Each zone’s median value of the fluorescence nano-sensor signal is used to create an optimal highly resolved local calibration.

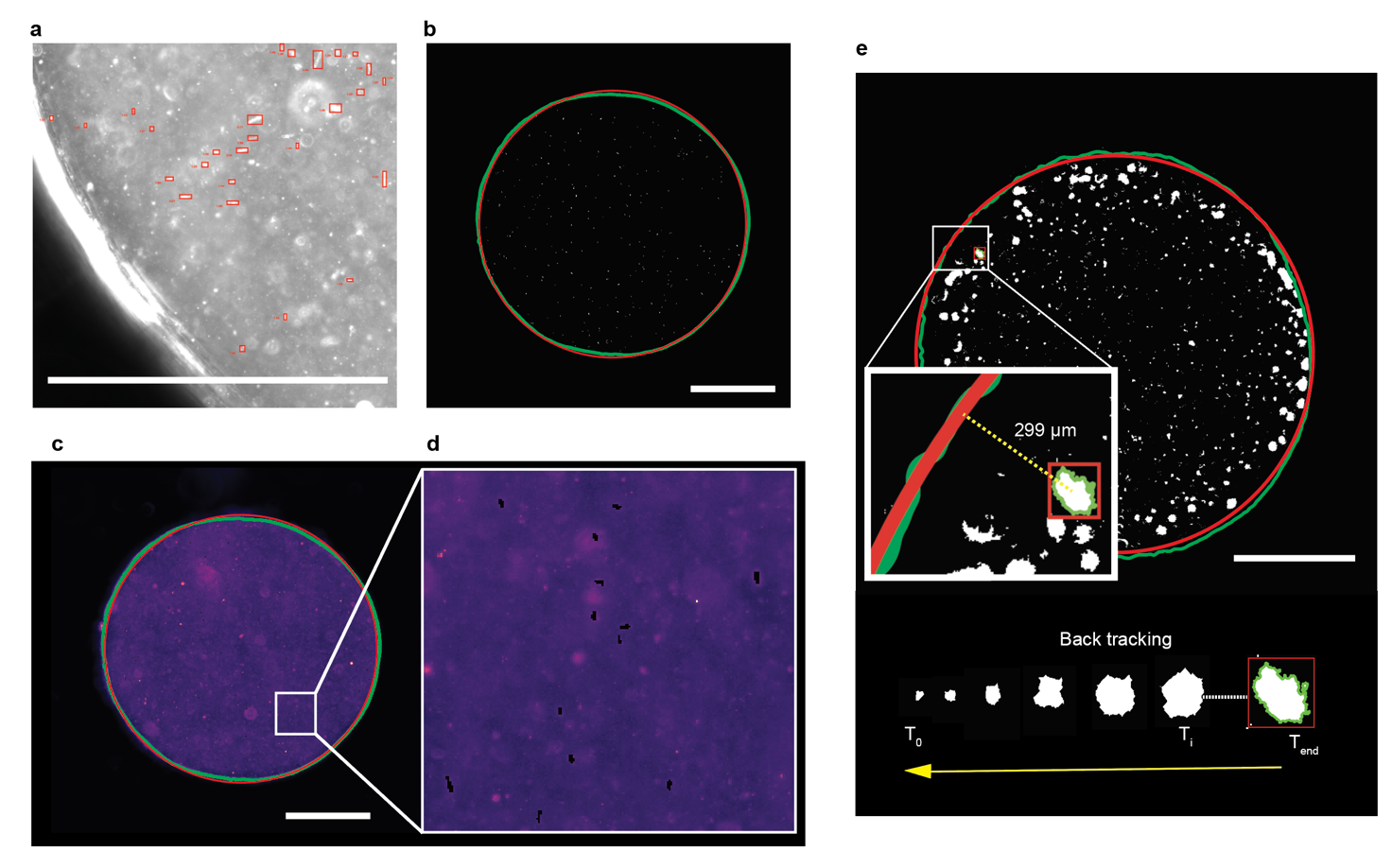

**Extended Data Fig. 4 Step-by-step image analysis algorithm. a)** Diatoms are recognized using an elongation shape factor. **b)** Binarized image of segmented diatoms. The edge of the hydrated particle is segmented and the contours are highlighted in green. In red is the fitted circle on the edge of the hydrated particle **c)** Oxygen nano-particles image across a full particle. The diatoms are subtracted from the fluorescence image to avoid interference by chlorophyll. **d)** Close up of diatoms subtracted from the fluorescence image. **e)** Each individual segmented colony is identified within a bounding box, and the distance from the edge of the hydrogel particle and area of the colony is recorded at each time point with a backtracking algorithm from the final measurement. The white scale bar in all panels is equal to 1000 µm.

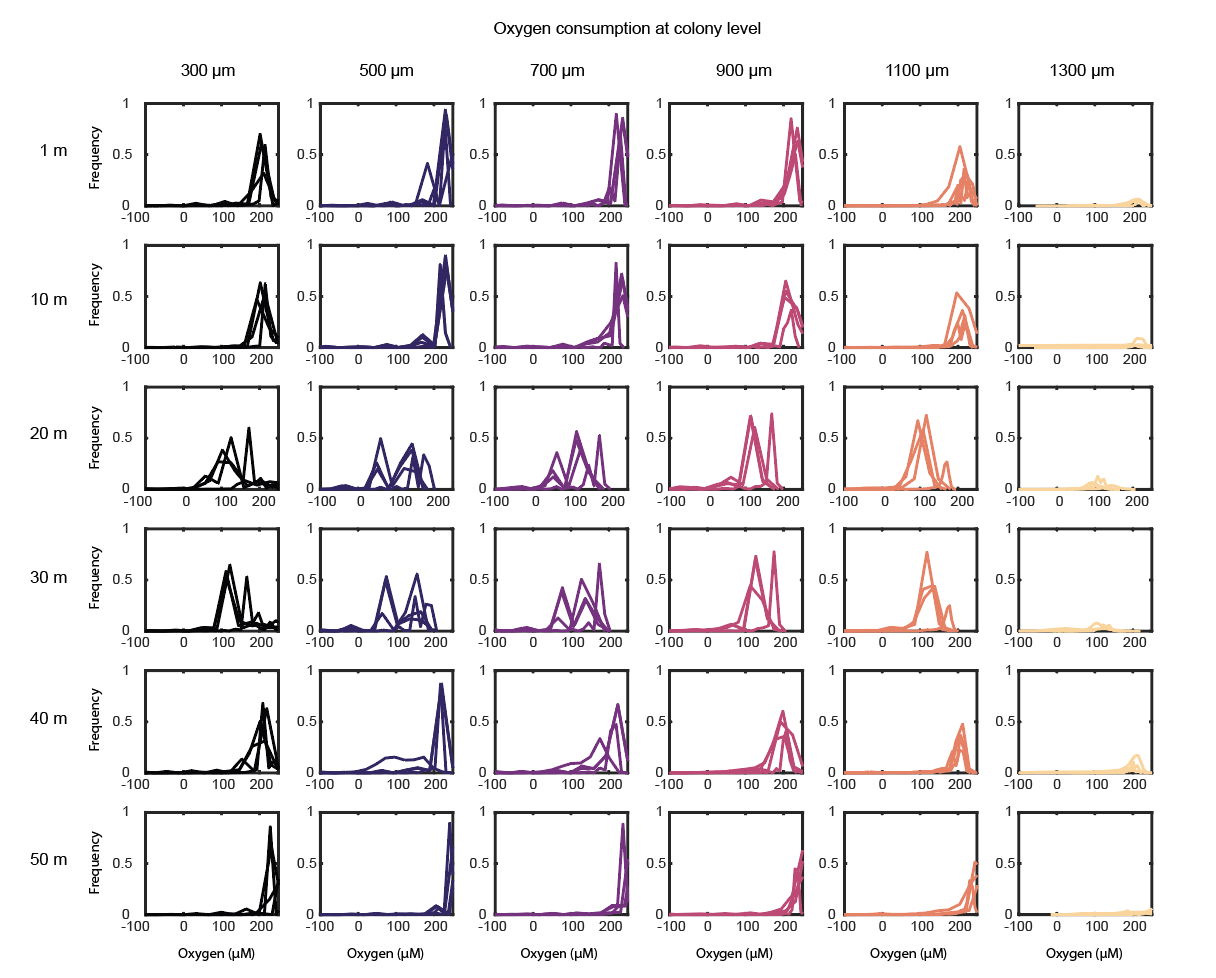

**Extended Data Fig. 5 Probability density function of oxygen change at colony level of particles seeded with broken diatoms.** At each time point (rows), the distribution is shown of the fluorescence signal of oxygen in all six spatial zones (columns) of all the colonies within each zone. Replicate particles n = 4.

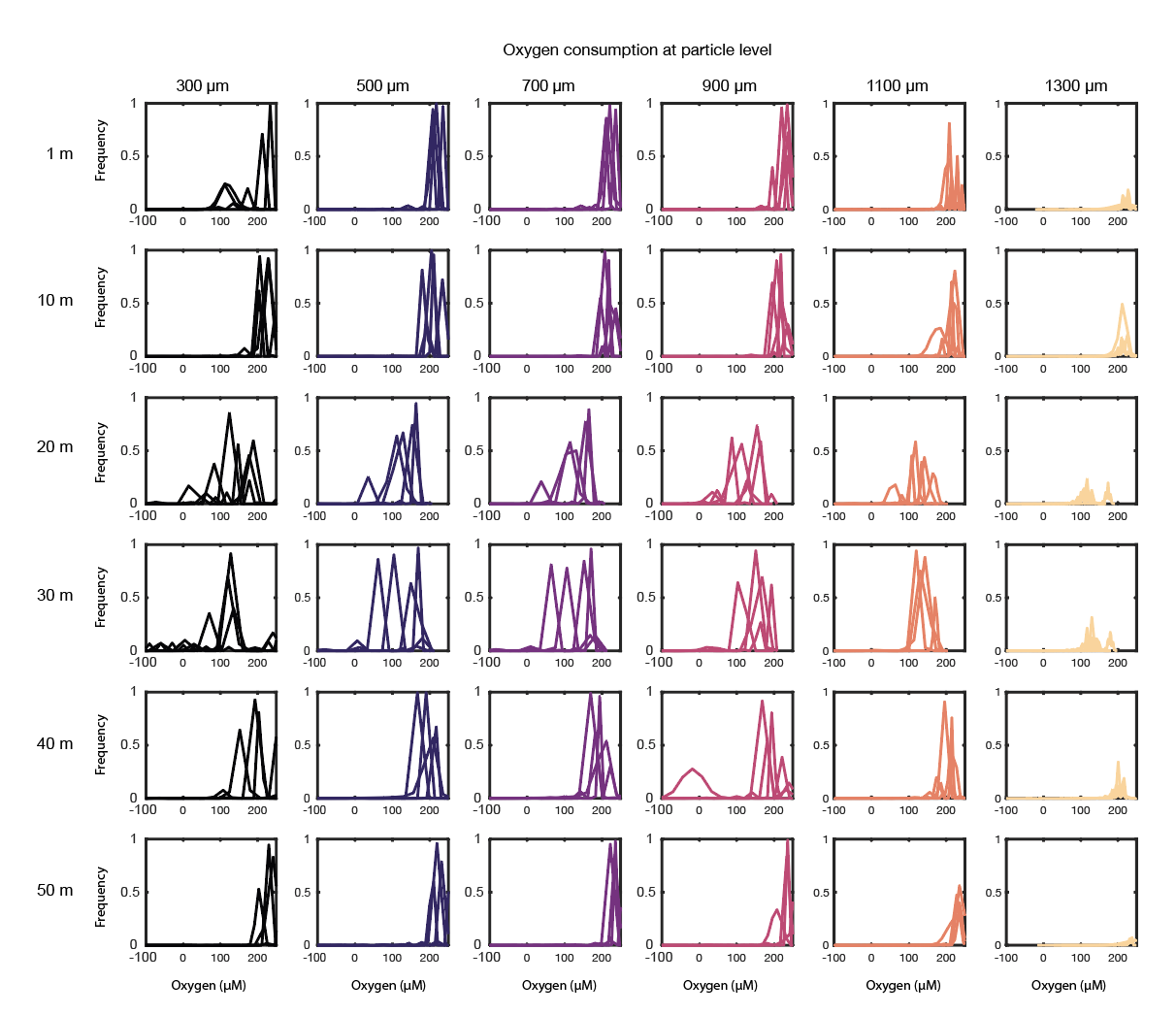

**Extended Data Fig. 6 Probability density function of oxygen change at particle level of particles seeded with broken diatoms.** At each time point (rows), the distribution is shown of the fluorescence signal of oxygen in all six spatial zones (columns) of the particle excluding the colonies. Replicate particles n = 4.

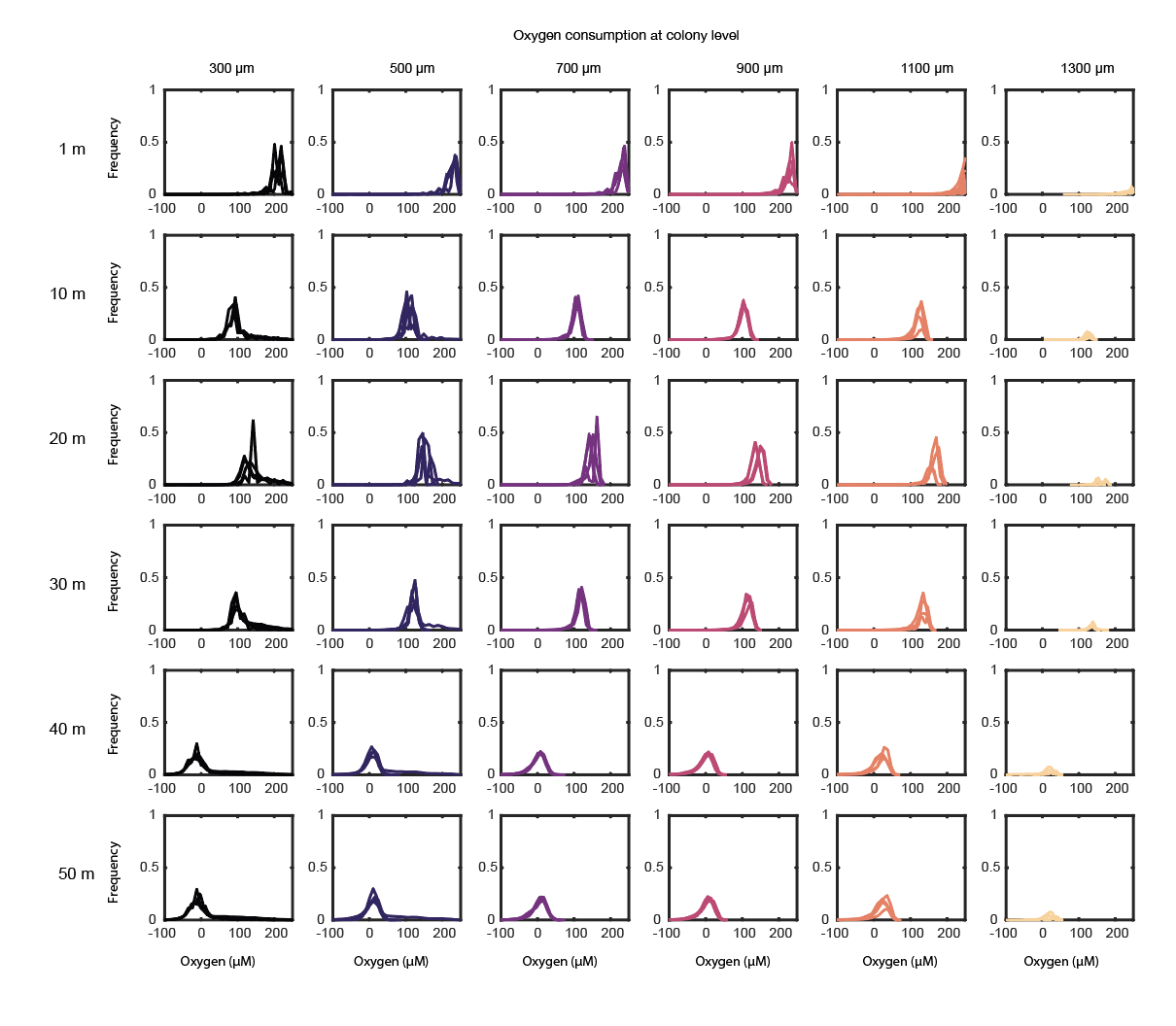

**Extended Data Fig. 7 Probability density function of oxygen change at colony level of particles seeded with intact diatoms.** At each time point (rows), the distribution is shown of the fluorescence signal of oxygen in all six spatial zones (columns) of all the colonies within each zone. Replicate particles n = 4.

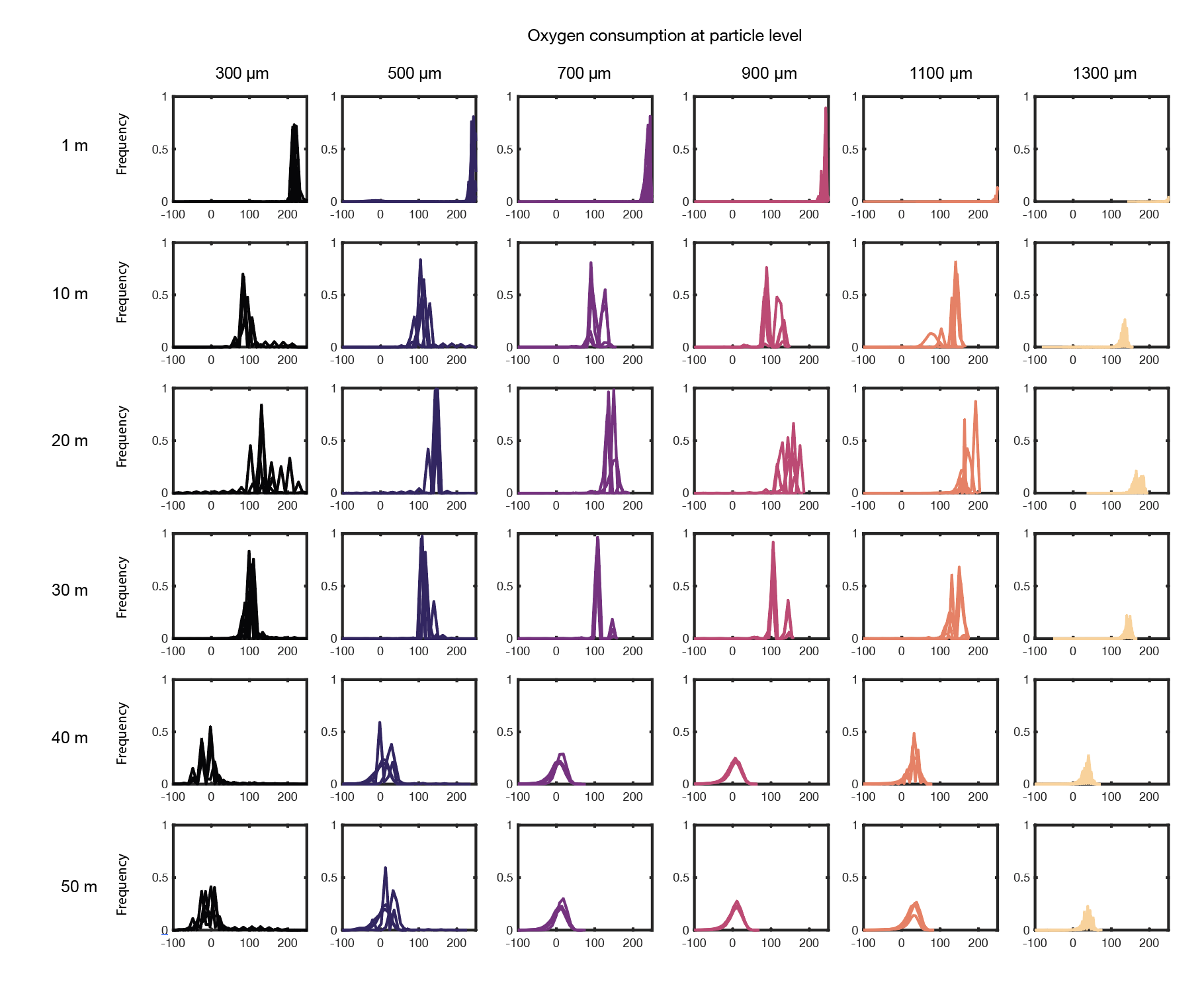

**Extended Data Fig. 8 Probability density function of oxygen change at the particle level of particles seeded with intact diatoms.** At each time point (rows), the distribution is shown of the fluorescence signal of oxygen in all six spatial zones (columns) of the particle excluding the colonies. Replicate particles n = 4.

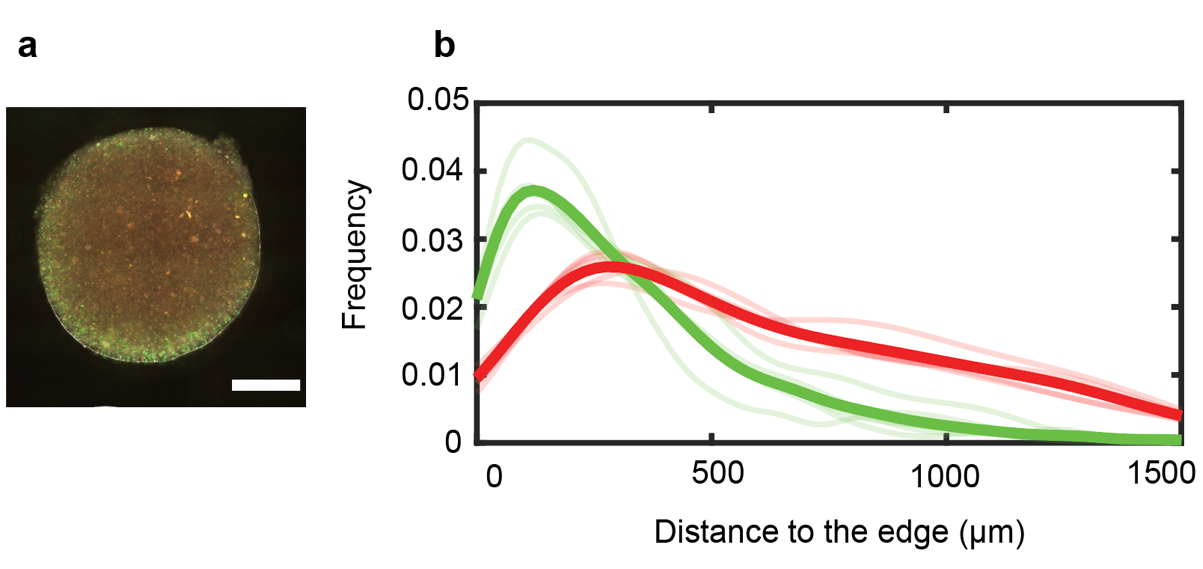

**Extended Data Fig. 9 Spatial expression of NarK and NirS genes for experiment with *Phaeodactylum tricornutum*.** a) Particle seeded with PAO1 NarK-GFP and PAO1 NirS-dsRed and *Phaeodactylum tricornutum*. White scale bar denotes 1000 µm. **b)** Probability density functions of the expression of NarK-GFP and NirS-dsRed reporters of five particles, with the average shown as the thick curve.

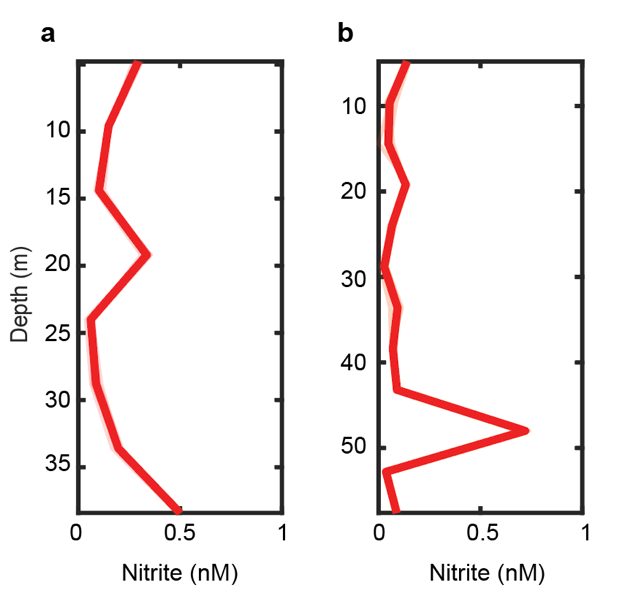

**Extended Data Fig. 10 Nitrite evolution in the water column of particles seeded with *Pseudomonas aeruginosa* and *Phaeodactylum tricornutum* or *Chaetoceros affinis*. a)** Nitrite measured in the bulk flowing seawater around 4 particles seeded with *P. aeruginosa* PAO1 NarK-GFP and PAO1 NirS-dsRed and *Phaeodactylum tricornutum*. 5 technical replicates are shown. **b)** Same as **a** but using *Chaetoceros affinis* as the diatom.

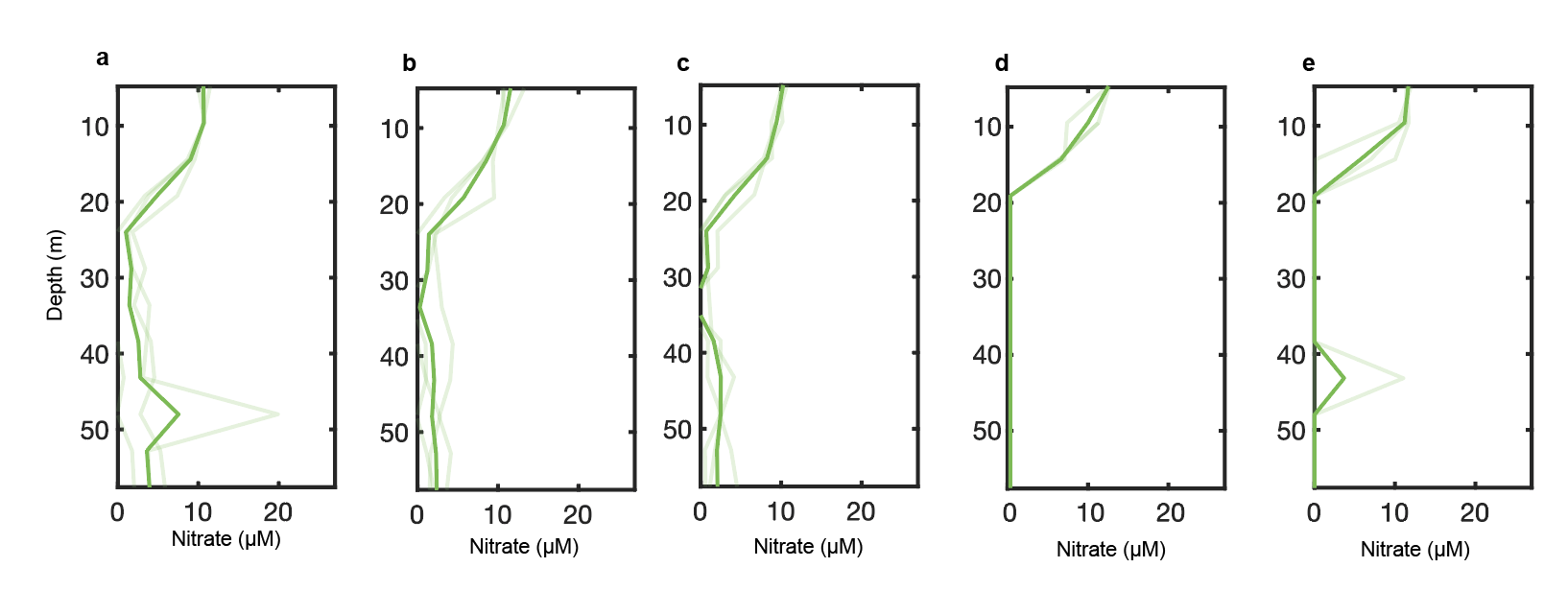

**Extended Data Fig. 11 Nitrate evolution in the water column.** Nitrate profiles measured in the water flowing around particles seeded with **a)** *Pseuomonas aeruginosa* PAO1 NarK-GFP and PAO1 NirS-dsRed and broken *Chaetoceros affinis*, **b)** broken *Chaetoceros affinis* carrying xenic bacteria, **c)** only with marine bacteria, **d)** *Pseudomonas aeruginosa* PAO1 NarK-GFP and PAO1 NirS-dsRed and intact *Chaetoceros affinis*, or **e)** intact *Chaetoceros affinis* carrying xenic bacteria. Three millifluidic devices containing 4 particles. Each panel shows 3 millifluidic device replicates, each containing 4 replicate particles.

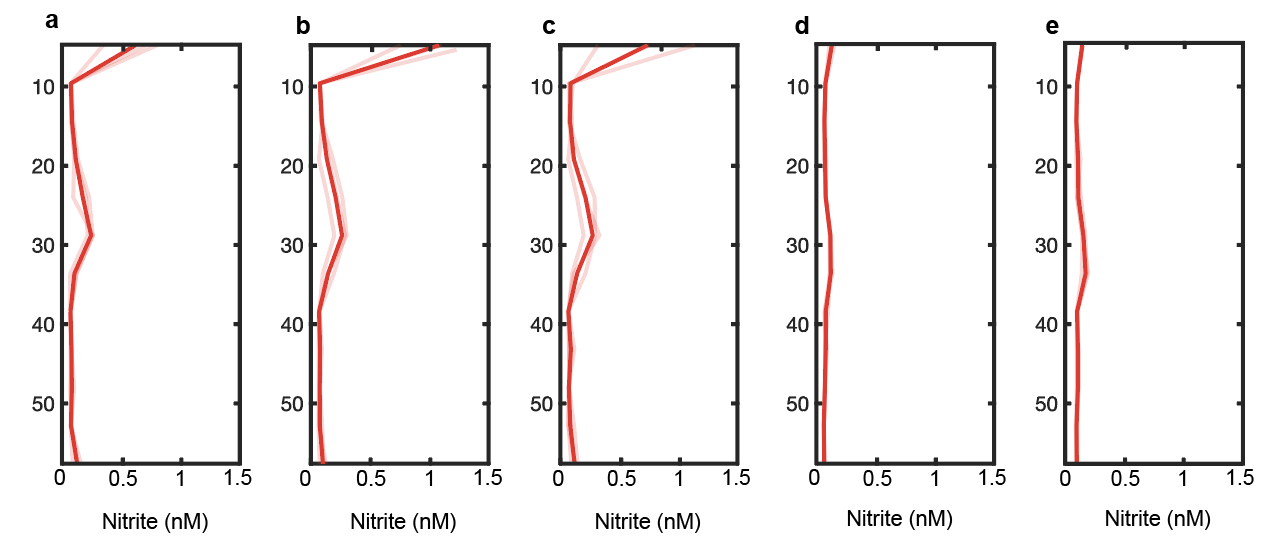

**Extended Data Fig. 12 Nitrite evolution in the water column.** Nitrite profiles measured in the water flowing around particles seeded with **a)** *Pseuomonas aeruginosa* PAO1 NarK-GFP and PAO1 NirS-dsRed and broken *Chaetoceros affinis*, **b)** broken *Chaetoceros affinis* carrying xenic bacteria, **c)** only with marine bacteria, **d)** *Pseudomonas aeruginosa* PAO1 NarK-GFP and PAO1 NirS-dsRed and intact *Chaetoceros affinis*, or **e)** intact *Chaetoceros affinis* carrying xenic bacteria. Three millifluidic devices containing 4 particles. Each panel shows 3 millifluidic device replicates, each containing 4 replicate particles.

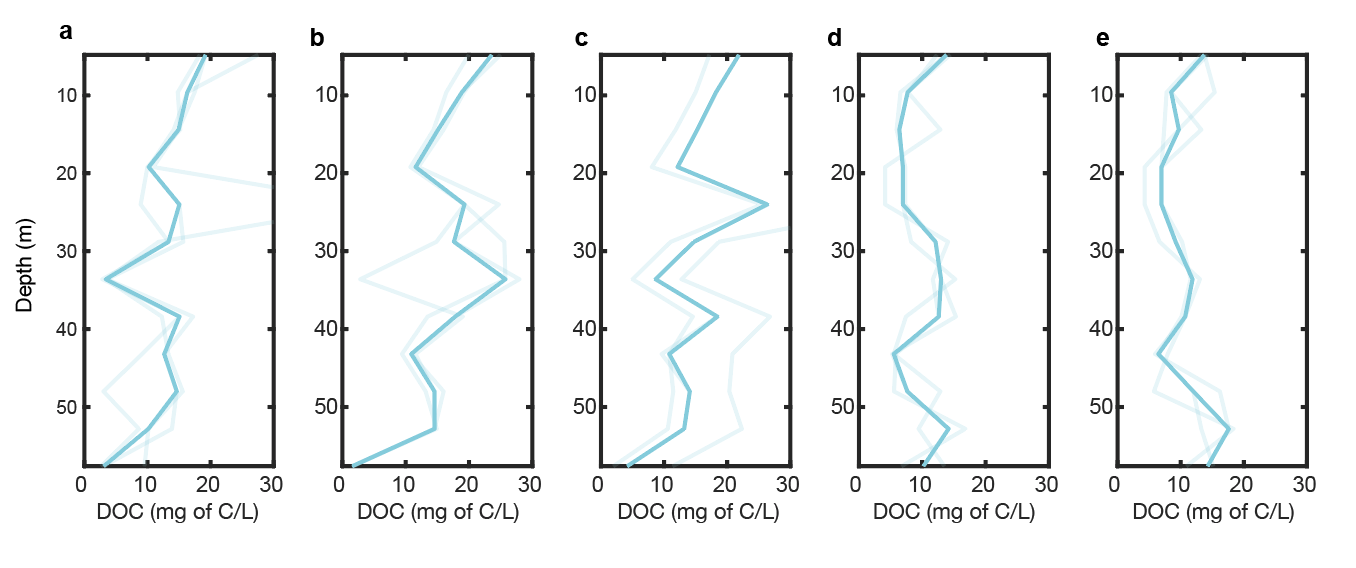

**Extended Data Fig. 13 DOC evolution in the water column.** DOC profiles measured in the water flowing around particles seeded with **a)** *Pseuomonas aeruginosa* PAO1 NarK-GFP and PAO1 NirS-dsRed and broken *Chaetoceros affinis*, **b)** broken *Chaetoceros affinis* carrying xenic bacteria, **c)** only with marine bacteria, **d)** *Pseudomonas aeruginosa* PAO1 NarK-GFP and PAO1 NirS-dsRed and intact *Chaetoceros affinis*, or **e)** intact *Chaetoceros affinis* carrying xenic bacteria. Three millifluidic devices containing 4 particles. Each panel shows 3 millifluidic device replicates, each containing 4 replicate particles.
